# Transient developmental increase in cortical projections to amygdala GABAergic neurons contribute to circuit dysfunction following early life stress

**DOI:** 10.1101/2022.04.21.489031

**Authors:** Joni Haikonen, Jonas Englund, Shyrley Paola Amarilla, Zoia Kharybina, Alexandra Shintyapina, Kristel Kegler, Marta Saez Garcia, Tsvetomira Atanasova, Tomi Taira, Henrike Hartung, Sari E. Lauri

## Abstract

Early life stress (ELS) results in enduring dysfunction of the cortico-limbic circuitry, underlying emotional and social behavior. However, the neurobiological mechanisms by which ELS affects development of the circuitry remain elusive. Here, we have combined viral tracing and electrophysiological techniques to study the effects of maternal separation (MS) on fronto-limbic connectivity and function in young (P14-21) rats. We report that aberrant prefrontal (mPFC) inputs to basolateral amygdala (BLA) GABAergic interneurons transiently increase the strength of feedforward inhibition in the BLA, which raises LTP induction threshold in MS treated male rats. The enhanced GABAergic activity after MS exposure associates with lower functional synchronization within prefrontal-amygdala networks *in vivo*. Intriguingly, no differences in these parameters were detected in females, which were also resistant to MS dependent changes in anxiety-like behaviors. Impaired plasticity and synchronization during the sensitive period of circuit refinement may contribute to long-lasting functional changes in the prefrontal-amygdaloid circuitry that predispose to neuropsychiatric conditions later on in life.

## Introduction

Cortico-limbic connectivity, underlying emotional behaviors, develops during an extended period of early life. During this time, the formation and refinement of the circuitry is particularly vulnerable to adverse experiences. Accordingly, early life stress (ELS) has been linked with increased risk for neuropsychiatric conditions involving emotional disturbance later on in life, both in humans as well as in rodent models (Tottenham, 2020).

Behavioral tests indicate that stress accelerates developmental emergence of emotional behaviors (reviewed by Callaghan et al., 2014; Ehrlich and Josselyn 2016; Bodegom et al., 2017), implying that ELS also facilitates maturation of the cortico-limbic brain circuits. In rodents, the cortico-amygdaloid projections appear during the second postnatal week, after which their functional maturation, involving strengthening of both pre- and postsynaptic signaling, goes on until approximately P30 (Bouwmeester et al., 2002, Arruda Carvalho et al., 2017). Consistent with the hypothesis of ELS accelerated development, data from longitudinal tracing studies suggest that ELS leads to precocious maturation of the amygdala-PFC connectivity, in a sex - specific manner (Honeycutt et al., 2020; Manzano-Nieves et al., 2020). However, the knowledge on the cellular and circuit mechanisms by which ELS influences this developmental process remains sparse. In particular, it is well established that projections from medial prefrontal cortex (mPFC) target both glutamatergic and GABAergic neurons in the adult basolateral amygdala (BLA) (Cho et al., 2013; Arruda-Carvalho and Clem, 2014; Hubner et al., 2014), yet no data exists on the effect of stress on the development of mPFC innervation to different cell types. Projections from the mPFC to the amygdala participate in the top-down control of emotional behaviors and shift in valence from positive to negative functional connectivity during development, in parallel with downregulation of amygdala reactivity (Gee et al., 2013). Developmental changes in the proportion of GABA vs. glutamatergic projection targets could critically influence the overall effect of cortical input on amygdala excitability and contribute to the developmental shift towards negative functional connectivity (Gee et al., 2013). Furthermore, since excitation/inhibition (E/I) balance strictly gates activity-dependent plasticity in the amygdala (Watanabe et al., 1995; Bissiere & Lüthi 2003; Shin et al., 2006; Bazelot et al., 2015), any alterations in the GABAergic activity during the critical period of circuit refinement would be expected to result in long-lasting changes in the connectivity.

Here we have used maternal separation (MS) as a model for ELS and studied its effects on development of mPFC projections to amygdala GABAergic neurons and on GABAergic transmission in 2-3 week old rats, the developmental stage when the cortical innervation to amygdala are emerging (Bouwmeester et al 2002; Arruda-Carvalho et al., 2017). Our data supports that MS increases projections from mPFC to amygdala GABAergic neurons, resulting in developmentally transient male-specific upregulation of GABAergic tone in the BLA. The increase in GABAergic activity after MS exposure associates with attenuated synaptic plasticity as well as impaired functional long-range interactions between mPFC and BLA in young male rats. Intriguingly, under our conditions, no differences in these parameters were detected in female rats, which were also resistant to ELS dependent changes in anxiety-like behaviors. Our data provides a novel mechanistic view and explains the discrepancy between increased anatomical but decreased functional connectivity data on the effects of ELS on cortico-amygdala networks.

## RESULTS

### Maternal separation (MS) increases anxiety-like behavior in male but not female rats

To validate that our protocol for maternal separation (MS) affected behavior later on in life, we performed two well-established tests for anxiety-like behavior: Open-field and elevated plus maze (EPM) tests. To this end, a total of 80 rats were divided into two groups with equal sex distribution. The pups in the MS group were separated from their dam for 3 hours daily during the postnatal days (P)2-14, while the littermate controls were reared under normal conditions. The behavioral testing was carried out at the age of 2 months (P60-P70).

The male MS rats showed significantly lower exploratory activity in the open field test as compared to the controls (Figure 1 A). In addition, the male MS rats travelled significantly less (percent of the total distance) in the central zone of the open field and in the open arms of the plus maze as compared to controls (Figure 1). However, no differences in these parameters were observed between female MS and control rats. These data indicate that MS produces behavioral changes consistent with anxiety-like phenotype in male but not female rats.

**Figure 1.**
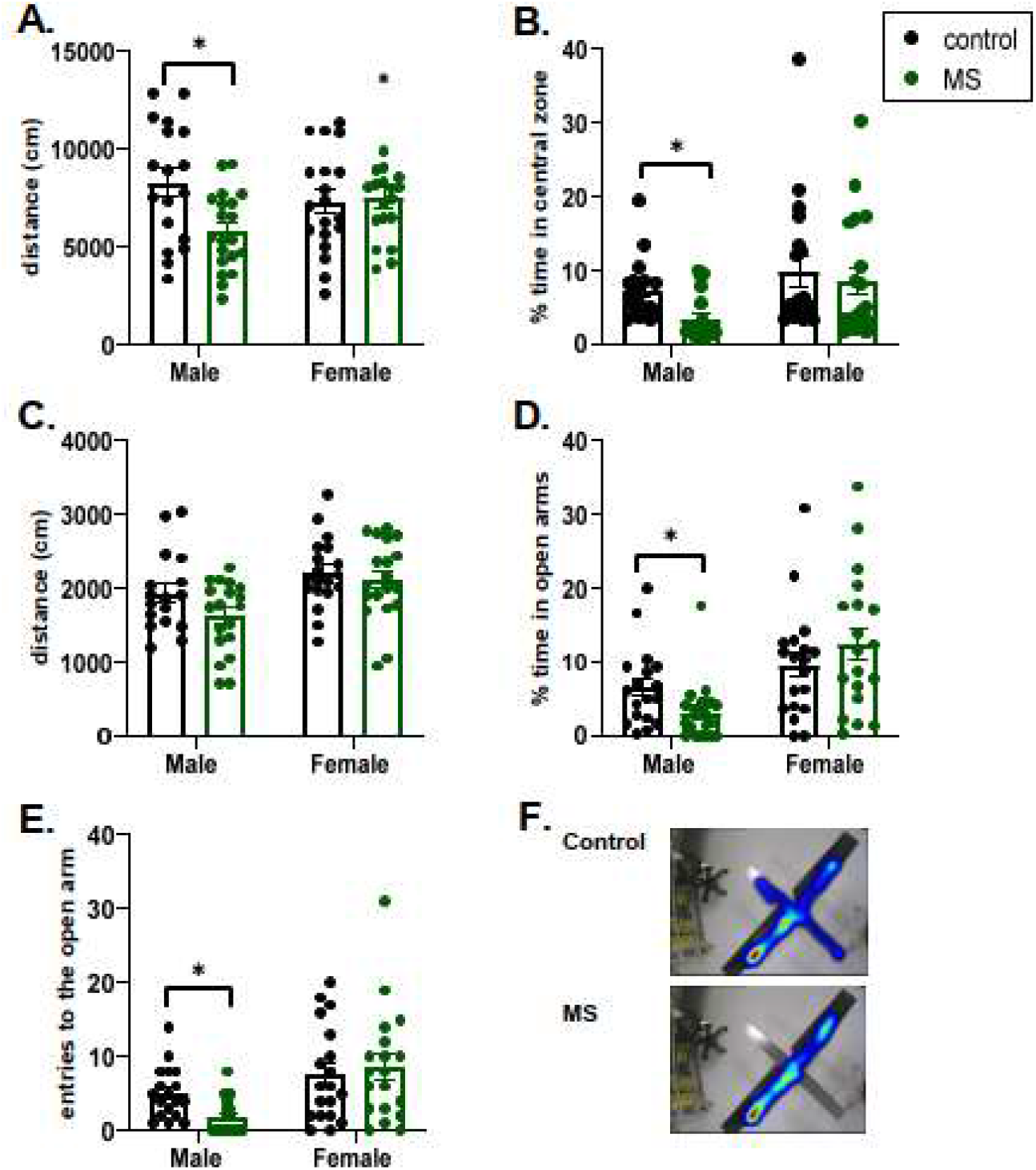
Maternal separation increases anxiety-like behavior in male rats. A. The total distance traveled in the open field arena (test time 1 h). There was a significant effect of MS on the total distance traveled (F_(1, 72)_ = 5.533, p=0.02) in males (control 8269 ± 698 cm, MS 5580 ± 430 cm; * p<0.003, Holm-Sidak) but not in females (control 7279 ± 580 cm, MS 7477 ± 506 cm). B. The % time spent in the central zone of the OF arena was significantly different between males and females (F_(1, 72)_ = 6.336, p=0.01) and affected by the MS treatment in males (control 7.3 ± 0.9 %, MS 3.5 ± 0.7 %, **p<0.005, one-way ANOVA on ranks) but not in females, control 9.9 ± 2 %, MS 8.6 ± 1.8 %). C. Total distance traveled in the EPM. There was a significant effect of sex, but not MS, on the total distance traveled (F_(1, 72)_ = 10.53, p=0.002). D. The % time spent in the open arms was significantly different between males and females (F_(1, 72)_ = 15.35, p=0.002) and affected by MS treatment in males (control 6.6 ± 1.2%, MS 3.2 ± 0.9%, *p<0.01, one-way ANOVA on ranks) but not in females (control 9.6 ± 1.7 %, MS 12.5 ± 2.1 %). E. Number of entries to the open arms in the same test was significantly different between males and females (F_(1, 72)_ = 13.35, p=0.005) and affected by MS treatment in males (control 5.1 ± 0.8, MS 1.9 ± 0.5, *p<0.001, one-way ANOVA on ranks) but not in females, control 7.5 ± 1.4, MS 8.5 ± 1.7). F. Heat map showing the activity in the EPM. Male control n=19, Male MS n=21; Female control n=20, Female MS n=20.

### MS affects development of cortical projections to BLA GABAergic neurons in a sex-specific manner

Recent studies have reported strengthening of the mPFC-BLA anatomical connectivity (Honeycutt et al., 2018; Manzano Nieves et al., 2020) and either strengthening or weakening of the functional connectivity following ELS (Johnson et al 2018; Guadagno et al., 2018; Bolton et al., 2018). To investigate the possibility that ELS differentially affects mPFC projections to BLA GABAergic and glutamatergic neurons, we used viral tracing combined with GAD67 immunohistochemistry in control vs. MS rats. Anterograde transsynaptic tracer (AAV1-EGFP) was injected into mPFC during the first postnatal week (P4-P6) during the maternal separation protocol, and the brains were fixed for histological analysis at P14-15 or at P21. For investigation of the adult connectivity, the injection was done at P30 and fixation at P50. A strong EGFP signal was regularly found at the injection site in the mPFC, covering both the IL and PL (mean area 0.90 ± 0.06 mm^2^ (P14-P21) and 1.12 ± 0.09 mm^2^ (P50), infection rate 2233 ± 150 EGFP^+^ neurons/ mm^2^ corresponding to 65 ± 3.1 % of DAPI^+^ cells; Figure 2A). Outside the injection area, strong EGFP labeling was detected in the BLA, the established target region of the mPFC projections (Figure 2B).

**Figure 2.**
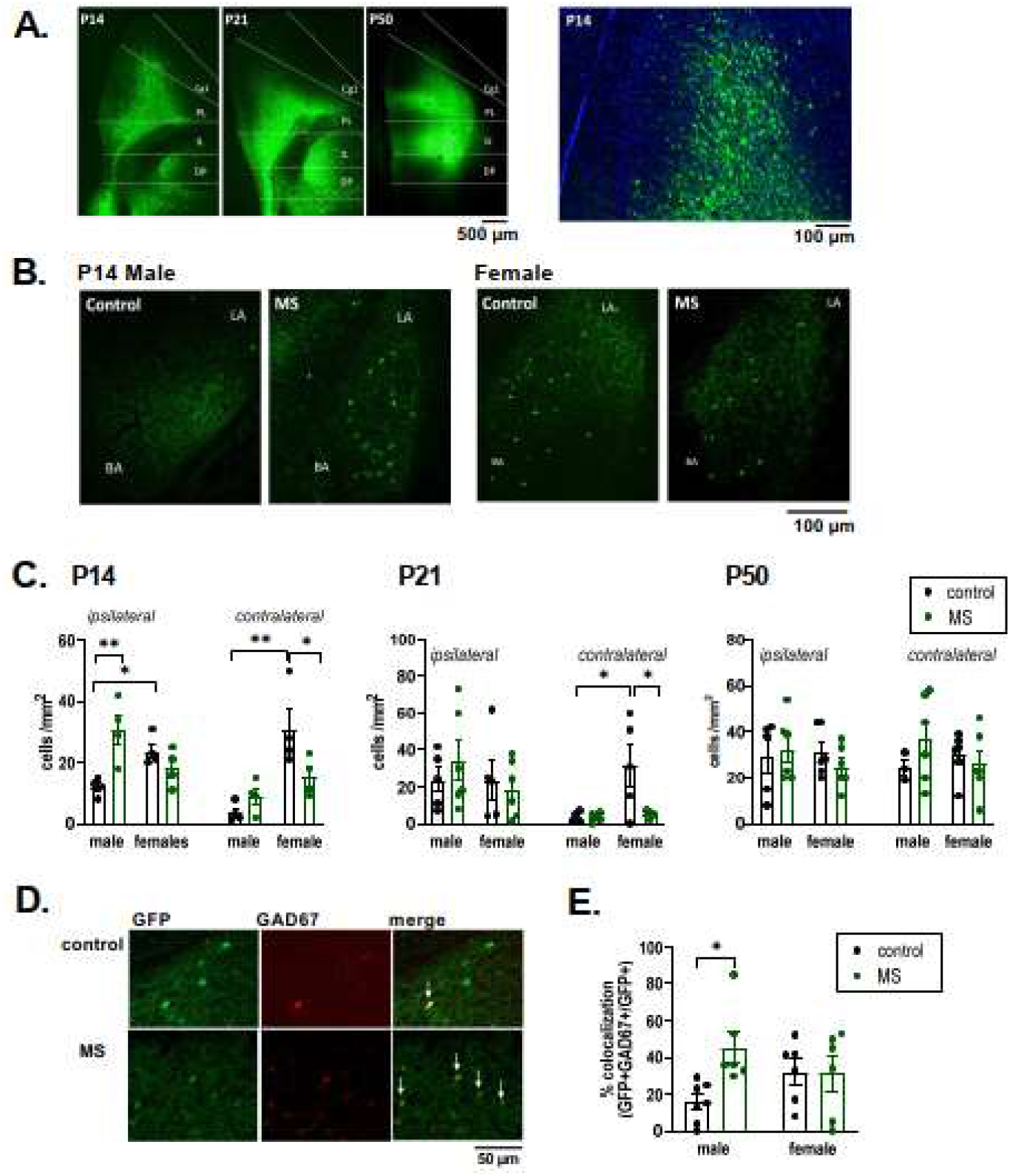
MS affects development of cortical projections to BLA GABAergic neurons. A. EGFP signal in the injection site (mPFC) at difference stages of development, 10-20 days after injection of the AAV1-GFP virus. Both prelimbic and infralimbic regions of the mPFC are strongly labeled with EGFP. The expanded image on the right shows the infected cells (65 ± 3.1 % of DAPI^+^ cells) at P14. PL: prelimbic; IL: infralimbic; Cg: cingulate cortex. B. EGFP expressing mPFC target neurons in the BLA. Example images from the brain of control and MS treated male and female rats. C. Quantification of the EGFP+ cell density in the BLA in control and MS treated male and female rats, at 3 different time points (P14, P21 and P50). There was a significant effect of sex on contralateral connectivity both at P14 (F_(1, 12)_ = 17.75,p=0.0012) and P21 (F_(1, 13)_ = 14.64, p=0.0021). At P14, the effect of MS depended on sex (ipsilateral, F_(1, 12)_ = 13.72,p=0.003; contralateral F_(1, 12)_ = 6.977, p=0.0215) and this effect persisted in the contralateral side at P21 (F_(1, 13)_ = 13.48, p=0.0028). Post-hoc tests revealed significantly higher connectivity in MS males (P14) in contrast to females showing lower connectivity as compared to controls at P14 and P21. Holm Sidak, *p<0.05; ** <0.001; ***p<0.0001. The individual dots represent averaged data from 8 sections / animal. P14 n=4/group; P21 n=5-6/group; P50 n=3-6/group. D. Example images showing co-localization (arrows) of EGFP and GAD67 staining (red) in the BLA, in sections from control and MS treated male rats. E. Quantified data for the percentage of GFP positive neurons that co-localize with GAD67 immunostaining in BLA (P14-21). The effect of MS on EGFP-GAD67 co-localization depended on sex (F (1, 21) = 4.483, p=0.0463) and was significantly higher in MS males as compared to controls (Holm-Sidak, *p= 0.015). n=6 / group

Quantification of the density of EGFP – positive neurons in the BLA in control rats indicated that at P14 but not later on in development, females have stronger anatomical connectivity between the mPFC and the BLA as compared to males in the hemisphere that was ipsilateral to the injection site (Figure 2B,C). Intriguingly, the sex differences in the connectivity were very pronounced on the contralateral hemisphere, both at P14 and P21 but disappeared towards adult stage (P50), apparently due to the delayed growth of contralateral projections in males (Figure 2C). At P14, 20 ± 3 % and 30 ± 9 % of the EGFP positive neurons in BLA colocalized with GAD67 immunostaining in males and females, respectively. The colocalization percentage was not significantly different between sexes (p=0.13, unpaired t-test) and did not change during development to P21 (male, 22 ± 5 % and female, 32 ± 5 %).

MS affected mPFC-BLA connectivity in a sex-specific manner. In P14 males, the density of EGFP^+^ neurons in the BLA was significantly higher in MS rats as compared to controls (Figure 2B,C). Similar trend was found at P21 and also in the contralateral hemisphere; however, these differences were smaller in magnitude as compared to P14 and did not reach statistical significance. In MS females, the density of EGFP^+^ neurons in the contralateral side was significantly lower as compared to controls both at P14 and P21, and a similar (non-significant) trend was observed on the ipsilateral side (Figure 2C). All these differences were developmentally transient and no longer observed at P50 (Figure 2C). Interestingly, the relative proportion of EGFP positive neurons that co-localized with GAD67 immunostaining was significantly increased by early life stress in males at P14-P21, while no differences were detected in females (Figure 2D,E).

These data show that the mPFC – BLA projections, particularly to the contralateral side, develop earlier in females as compared to males. Furthermore, higher mPFC – BLA connectivity in the MS vs. control males is consistent with accelerated maturation of the mPFC – BLA projections following ELS. Finally, increase in the percentage of EGFP-GAD67 colocalization suggests that early life stress specifically facilitates development of cortical projections to GABAergic interneurons in males.

### MS is associated with acute sex-specific changes in glutamate-driven GABAergic synaptic activity in the LA

Developmental changes in the proportion of excitatory inputs to GABAergic vs. glutamatergic neurons in the BLA could critically influence amygdala excitability. In order to investigate the effects of MS on E/I balance in the amygdala, we performed electrophysiological recordings from BLA neurons in acute slices from 14-15 day old (P14-15) and adult (P50-P80) control and MS rats. Whole cell voltage clamp recordings of spontaneous GABAergic (sIPSCs) and glutamatergic (sEPSCs) synaptic currents were carried out under experimental conditions were GABAergic events were seen as outward currents and glutamatergic events as inward currents (Figure 3A). No significant differences in the frequency or amplitude of sEPSC were detected between control and MS rats in lateral amygdala (LA) or basal amygdala (BA), in either males or females (Figure 3B; see Supplementary Figure 1 for amplitude data). However, in LA, sIPSC frequency was significantly higher in MS male rats as compared to controls (Figure 3C), in contrast to females where sIPSC frequency was lower in MS rats. In BA neurons, no difference in sIPSC frequency was detected between control and MS groups. Interestingly, both in LA and BA the basal frequency of sIPSCs was significantly higher in females as compared to males (Figure 3C, values). These differences were developmentally transient and were not observed in the adult stage (Figure 3D).

**Figure 3.**
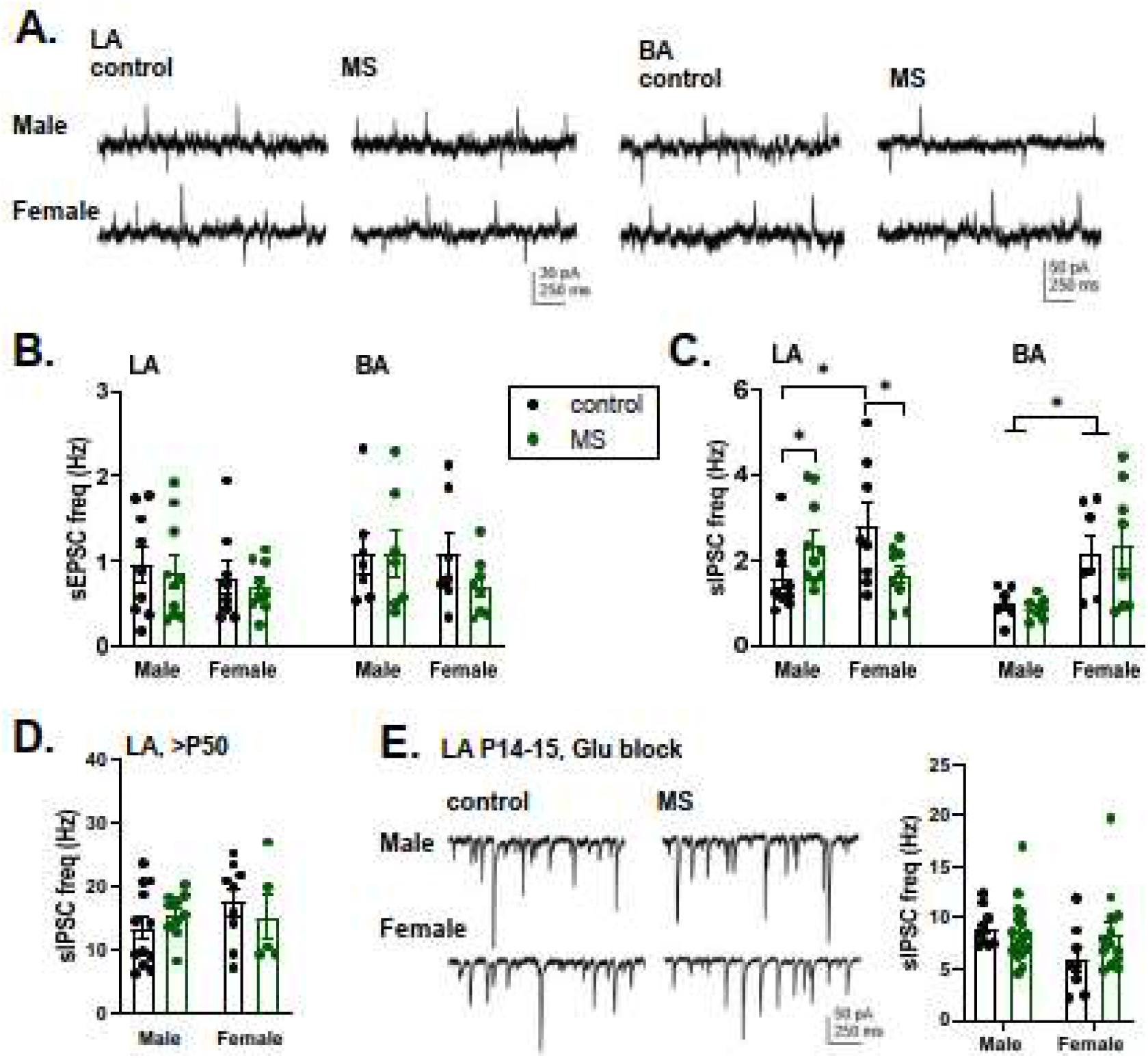
MS is associated with developmentally restricted and sex specific changes in GABAergic activity in the amygdala. A. Example traces depicting glutamatergic (inward currents) and GABAergic (outward currents) synaptic events in BA and LA neurons in slices from control and MS treated juvenile (P14-15) rats. B. Pooled data on the frequency of sEPSCs in BA and LA neurons in recordings from control (black symbols) and MS (green symbols) juvenile (P14-15) rats [control male n= 9(4 animals), female n=8(3), MS male n=10 (4), female n=9(3)]. No significant differences in sEPSC frequency between groups were detected. C. Pooled data on the frequency of sIPSCs, from the same cells as in B. The effect of MS on sIPSC frequency was significantly dependent on sex in LA (F_(1, 32)_ = 8.100, P=0.0077) and there was a significant effect of sex in BA (F_(1, 25)_ = 14.12, P=0.0009). In LA, MS treatment increases sIPSC frequency in males, decreases it in females and sIPSC frequency is higher in control females vs. males. (* p<0.05; one –way ANOVA on ranks). D. The frequency of sIPSC in LA neurons from adult control and MS treated rats [control male n=12(10), female n=9(7), MS male n=10 (8), MS female n=5(3)]. No significant differences between groups were detected. E. Example traces and pooled data on the frequency of sIPSCs in LA neurons from juvenile (P14-15) control and MS rats, recorded in the presence of CNQX to block glutamatergic activity. These recordings were done using high chloride concentration in the pipette filling solution, which improves detection of small GABAergic events that are now observed as inward currents. Under these conditions, no significant differences in sIPSCs were detected between the groups [control male, n=9(3), female n=8(3); MS male, n=16 (4), female n=13(4)].

These data indicate that MS treatment transiently influences GABAergic transmission in the LA, but has no apparent effects on glutamatergic synaptic transmission. The sex-specific changes in GABAergic transmission between control and MS rats were similar to the observed differences in mPFC –BLA connectivity, which mediate glutamatergic excitation of BLA interneurons (Hubner et al., 2014; Lucas et al., 2016). To study whether the altered GABAergic transmission in young MS rats depended on fast excitatory synaptic activity, we recorded sIPSCs in the presence of CNQX and AP5, antagonists of AMPA/KA and NMDA types of ionotropic glutamate receptors. Under these conditions, no differences in sIPSC frequency or amplitude were detected between control and MS groups (Figure 3E).

ELS-dependent changes in density and maturation of GABAergic interneurons have been described in adolescent rodents (e.g. Giachino et al., 2007; Seidel et al., 2008; Santiago et al., 2018; Manzano Nieves et al., 2020) and could contribute to the observed alterations in GABAergic transmission in MS rats. To address this possibility, we used qPCR and immunohistological stainings to study expression of marker proteins related to maturation of GABAergic neurotransmission in the BLA of control vs MS rats. qPCR analysis indicated no significant differences in the expression of mRNAs encoding for parvalbumin (*Parv*), somatostatin (*Sst*), KCC2 (*Slc12a5*) and NKCC1 (*Slc12a2*) in the BLA between control and MS rats, neither at P14 or P50 (Supplementary Figure 2A). Immunohistological staining indicated that the density of parvalbumin (PV) expressing GABAergic neurons was significantly different between sexes (p=0.02, two way ANOVA) and significantly lower in MS group as compared to controls (p=0.003, two way ANOVA). In contrast, there were no differences in the density of somatostatin (SST) expressing neurons in the BLA (Supplementary Figure 2B). Lower density of PV cells is unlikely to explain the observed increase in the GABAergic transmission in MS males. Together with the finding that the differences in sIPSC frequency were dependent on intact glutamatergic transmission, these data suggest that changes in glutamatergic drive to interneurons underlie the MS associated changes in the GABAergic activity in the juvenile amygdala.

### MS enhances the strength of feedforward inhibition in the BLA in response to activation of prefrontal inputs in males

Both LA and BA receive projections from various regions of the brain, all contributing to spontaneous synaptic inputs. To study the specific role of mPFC inputs on E/I balance in the BLA, we used optogenetic excitation of mPFC axons and studied the strength of feedforward inhibition (FFI) within LA and BA microcircuits. Injection of AAV virus encoding for EYFP-ChR2 under the CaMKII promoter into mPFC at P4-7 resulted in expression of EYFP in 37 ± 1.4 % of DAPI^+^ cells at P18-21 and the expression covered both PL and IL regions (Figure 4A). Light-evoked disynaptic IPSC (dIPSC)/EPSC ratio was measured in LA and BA principal neurons at P18-21 by recording a monosynaptic EPSC at −70 mV, after which the cell was clamped to 0 mV to isolate the dIPSC, mediated via activation of feedforward inhibitory interneurons.

**Figure 4.**
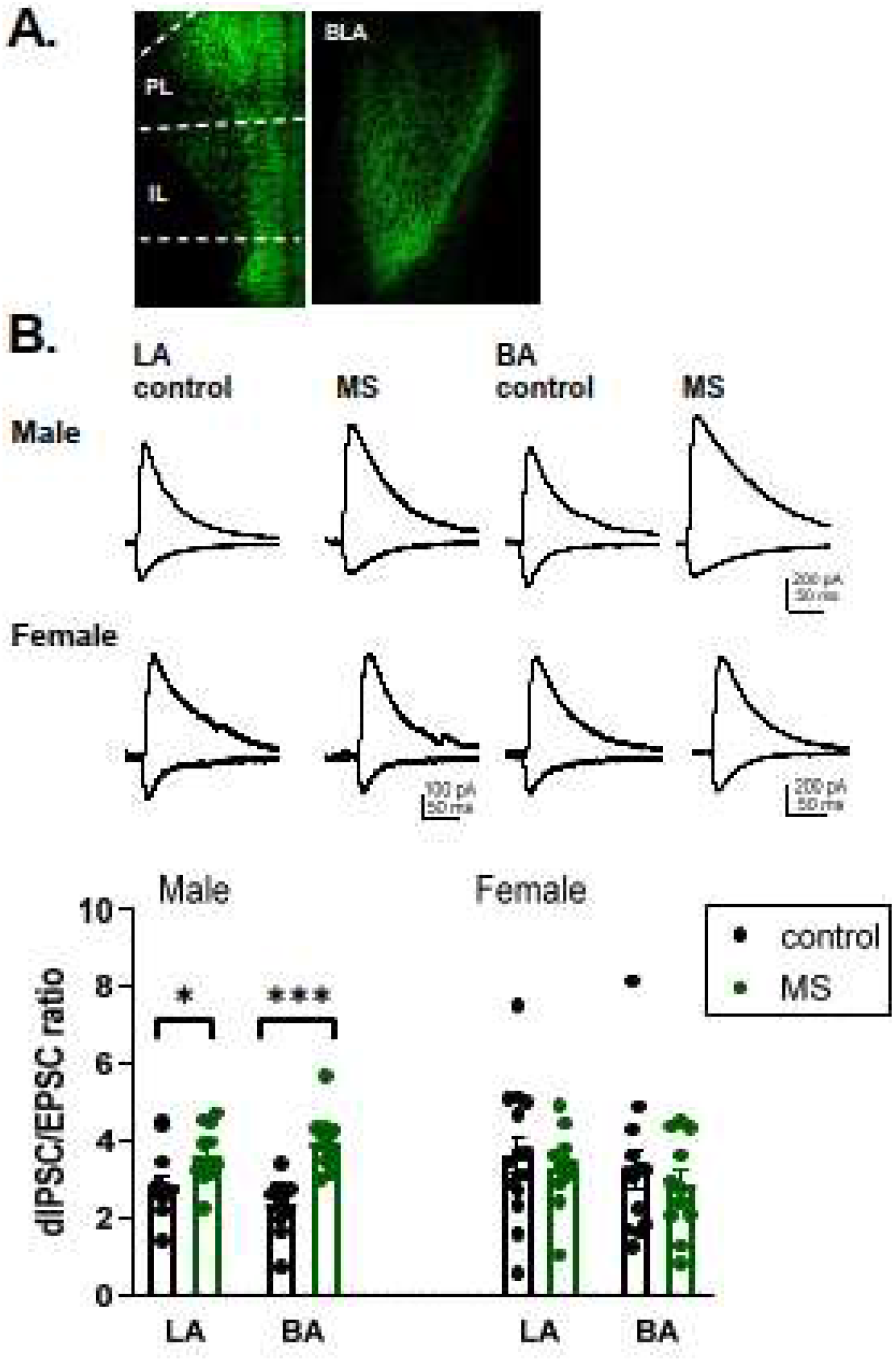
MS increases feedforward inhibition in the cortico-amygdaloid circuit in males. A. Expression of channelrhodopsin-EYFP in the mPFC at P18, two weeks after viral transduction (AAV-CaMKIIa-hChR2-EYFP). EYFP expression was observed in 37.23 ± 3.63 % of DAPI positive cells in both prelimbic and infralimbic regions of the mPFC. EYFP labeled axons were clearly visible in the BLA. B. Example traces and pooled data on the strength of feedforward inhibition (dIPSC/EPSC ratio) in LA and BA principal neurons, in response to optogenetic stimulation of mPFC inputs. dIPSC/EPSC ratio was significantly higher in MS males as compared to controls (F_(1,44)_ = 30.36, p<0.0001), in both LA (p=0.04, Holm-Sidak) and BA (p<0.0001, Holm-Sidak). No significant differences were detected in females. Male control, n= 13(4) for both LA and BA; Male MS n= 11 (4) for both LA and BA. Female control n=15(5) (LA) and 13(4) (BA), Female MS n=11 for both LA and BA. Pooled data represent average ± SEM.

In control rats, the dIPSC/EPSC ratio was higher in females as compared to males (F_(1,50)_ = 4,797, p=0.03), while there were no significant differences between LA and BA. MS treatment significantly increased the dIPSC/EPSC ratio both in LA and BA in males but had no effect in females (Figure 4B).

These data are consistent with the idea that in the BLA of male but not female rats, MS accelerates the development of the circuitry responsible for FFI in response to activation of prefrontal inputs. The increase in the dIPSC/EPSC ratio in MS male rats can be fully explained by an increase in excitatory projections from mPFC to BLA GABAergic neurons.

### Strong GABAergic activity in the BLA increases the threshold for LTP induction in MS males

GABAergic inhibition represents a key mechanism regulating induction of associative synaptic plasticity in the amygdala (Watanabe et al., 1995; Bissiere et al., 2003; Shin et al., 2006; Bazelot et al., 2015). To study whether the enhanced FFI in male MS rats correlated with changes in synaptic plasticity, we studied LTP in the cortico-amygdaloid pathway *in vitro*. Electrical stimulation of the external capsulae was used to mimic activation of cortical afferents in acute slices from P17-18 control and MS rats, and glutamatergic postsynaptic field responses (fEPSPs) were recorded from the BLA. Theta burst afferent stimulation (TBS) produced a mild but repeatable LTP of fEPSPs in male control rats (Figure 5A). TBS induced synaptic potentiation was significantly smaller in slices from male MS rat pups as compared to controls (Figure 5A). Interestingly, in female rats of corresponding age, TBS produced no significant potentiation in either control or MS groups (Figure 5B).

**Figure 5.**
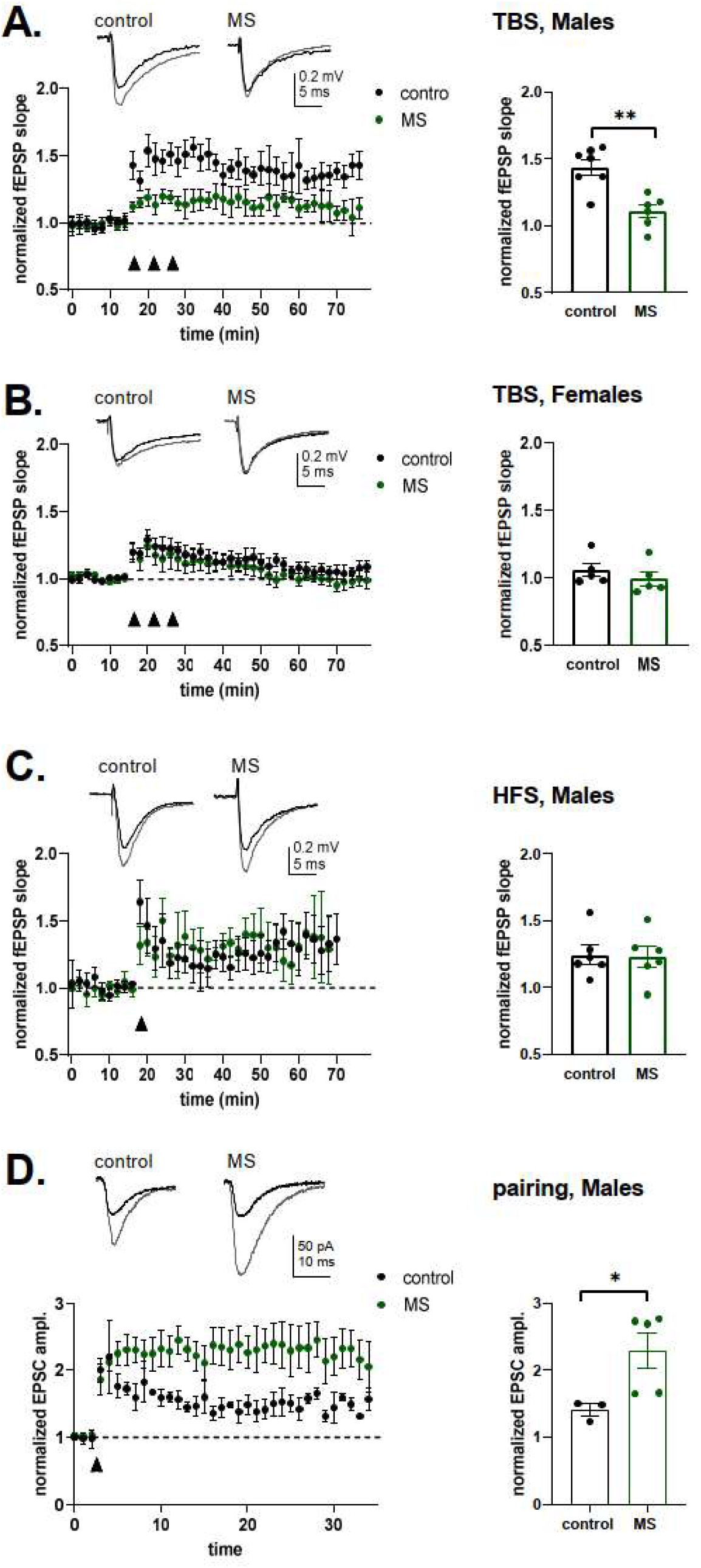
MS is associated with an increase in the LTP induction threshold in the cortico-amygdaloid pathway in males. A. A time course plot showing the effect of TBS (arrowheads) on fEPSPs, recorded from BLA in response to stimulation of cortical afferents in slices from control and MS exposed male rats. Superimposed example traces before (black) and 60 min after (grey) TBS are shown on top. Pooled data shows the average level of synaptic potentiation 60 minutes after the TBS in acute slices from control (n=7(3)) and MS (n=6(4)) male rats. The level of synaptic potentiation was significantly lower in the MS group (p=0.0012, unpaired two tailed t-test). B. Similar data as in A, for control (n=5(3)) and MS females (n=5(3)). No significant differences were observed between the groups. C. Similar data as in A, for the effect of HFS (100Hz, 1s) on fEPSPs in control (n=6(4)) and MS (n=4(2)) males. D. Example traces and pooled data showing LTP induction in the presence of GABA-A receptor antagonist picrotoxin, in acute slices from control and MS male rats. The data is collected in whole cell voltage-clamp mode. Pairing membrane depolarization (+10 mV) to theta-burst afferent stimulation produces a robust LTP in both control (n=3(2)) and MS (n=5(3)) groups. The average level of synaptic potentiation was significantly higher in the MS group 30 min after the pairing (p=0.047, unpaired two tailed t-test).

A stronger stimulation protocol (high –frequency tetanic stimulation, HFS) produced LTP in both control and MS males (Figure 5C), indicating that in the MS rats, there was not a deficiency to express LTP *per se*, but rather an increase in the threshold for its induction. Accordingly, a robust LTP was induced both in control and MS rats when using a pairing protocol in the presence of GABA-A receptor inhibitor picrotoxin (Figure 5D). Interestingly, the level of pairing induced LTP was higher in the MS groups as compared to controls. Together, these data indicate that the heightened inhibitory drive associated with early life stress increases threshold for induction of LTP in the BLA.

### MS impairs functional coupling between prelimbic (PL) cortex and BLA in males

The increased anxiety-like behaviour, the abnormal development of prefrontal-amygdala anatomical connectivity as well as the increased FFI in the amygdala after ELS suggests that also prefrontal-amygdala functional interactions may be affected by ELS.

To test this hypothesis, simultaneous multi-site recordings of local field potentials (LFP) and multi-unit activity (MUA) were performed from PL and BLA of lightly urethane anaesthetized P14-15 male control (n=11), female control (n=14), male MS (n=13) or female MS (n=13) pups, respectively. Network activity in both regions was already continuous at this age and consisted of the previously described sleep-like rhythms mimicked by urethane anesthesia (Clement et al., 2008; Pagliardini et al., 2013). Large-amplitude slow oscillations (0.3-0.5 Hz), which were more prominent in PL than BLA, were superimposed by faster rhythms in delta/ low theta (2-5 Hz) and gamma (20-50 Hz) range (Fig. 6, A (iii), B(iii), Suppl. Fig 3). We focused our analysis on activity between 2-5 Hz, since activity in delta and low theta range has previously been identified as a substrate of functional communication between mPFC and BLA during expression of learned fear but also innate anxiety (Likhtik et al., 2014; Harris and Gordon, 2015; Dejean et al., 2016; Karalis et al., 2016). The power in this band was similar in male and female pups (Suppl. Fig 3, C (i), D (i)) and was not affected by ELS exposure (Suppl. Fig 3 A, B). To investigate if the coupling by synchrony between prefrontal-amygdala networks is altered by ELS exposure, we analyzed phase-phase coupling by computing the phase locking value (PLV) between LFP in PL and BLA in all four groups of pups for frequencies from 1-100 Hz excluding the slow anesthesia induced rhythms. Strongest phase-phase coupling in control rats was observed in a peak between 2-5 Hz that was significantly larger than phase-phase coupling calculated for shuffled data and similar in both male and female control pups (Fig. 6 D. (i)). Generally, when comparing the groups by Two-way ANOVA with factors treatment and sex, we did not find a significant effect of treatment, sex nor interaction. However, in male pups exposed to MS, PLV between 2-5 Hz was significantly decreased compared to controls as revealed by Holm-Sidak post hoc test (p=0.036, Fig. 6 C (i)). This effect of MS on prefrontal-amygdala coupling was specific for males since PLV in the same band was not different between control and MS exposed pups in females (Fig. 6 C (ii)). Finally, when comparing PLV between MS exposed males and females with Holm-Sidak post hoc test (Fig. 6 D (ii)), males had significantly lower (p=0.023) 2-5 Hz PLV than females.

**Figure 6.**
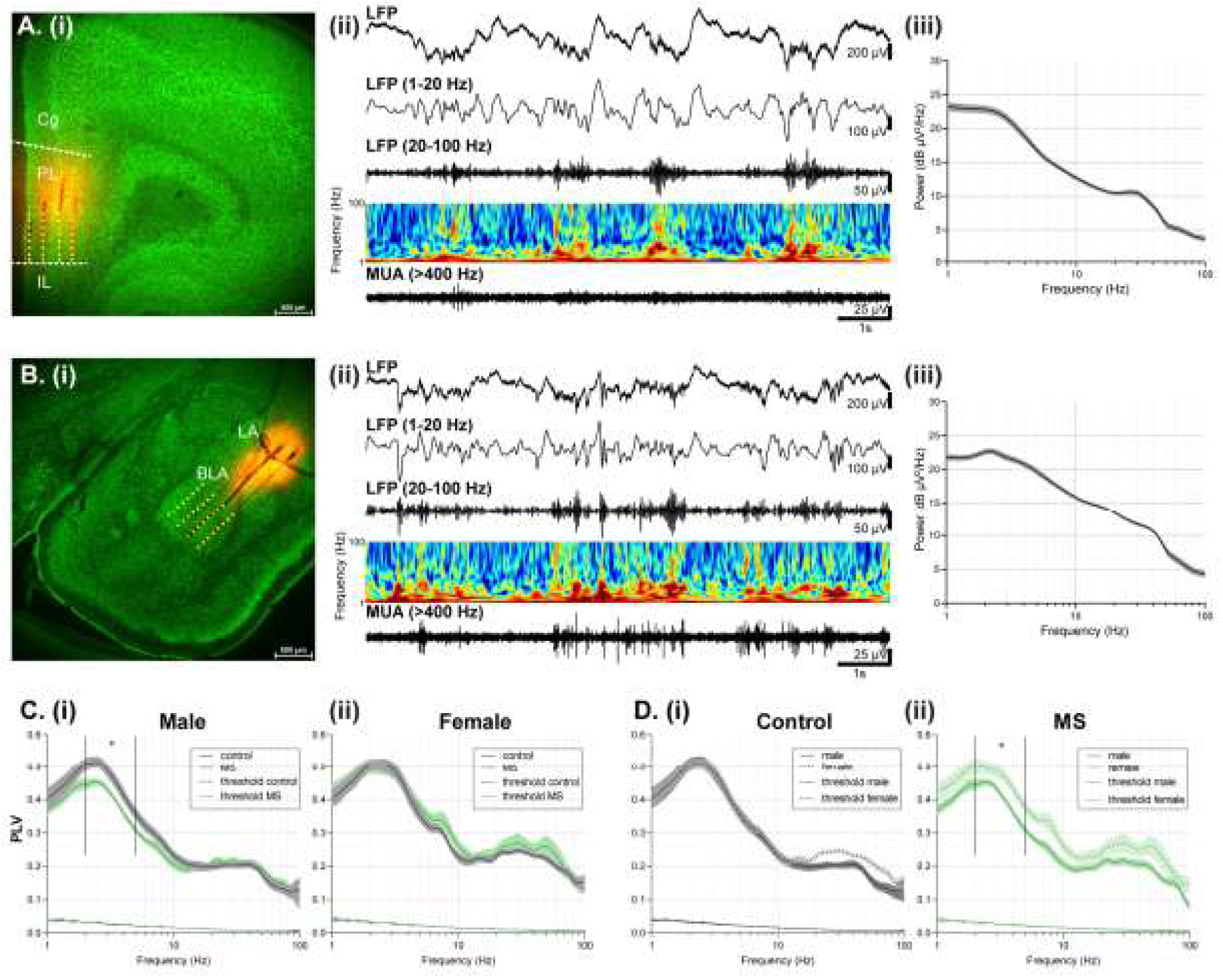
MS decreases prefrontal-amygdala oscillatory coupling in male pre-juvenile rats only. A. (i) Digital photomontage showing the location of the DiI-labelled 4-shank recording electrode (orange) in PFC of a 100 μm-thick coronal section from a P15 rat pup. The superimposed yellow dots mark the 32 recording sites (8 per shank) covering superficial and deep layers of PL. (ii) Extracellular LFP recording of oscillatory activity in PL from a P15 rat pup displayed after band-pass filtering (1-20 Hz, 20-100 Hz) and after 400 Hz high-pass filtering showing corresponding MUA activity. At this age, oscillatory activity is continuous with predominant slow oscillations and superimposed activity in delta, theta and gamma range. Color-coded frequency plots show the wavelet spectra at identical time-scale. (iii) Average logarithmic power-spectrum of prelimbic LFP of male control pups (n=11) showing broad peaks between 2-5 Hz and 20-50 Hz. B. (i) Digital photomontage showing the location of the DiI-labelled 4-shank recording electrode (orange) in amygdala of a 100 μm-thick coronal section from a P14 rat pup. The superimposed yellow dots mark the 32 recording sites (8 per shank) covering BLA. (ii) Same as A (ii) for corresponding oscillatory and MUA activity in BLA. (iii) Same as A (iii) for amygdala LFP of male control pups. C. (i) Average PLV spectra between activity in PL and BLA of male control (n=11, black) and MS pups (n=13, green) for original (straight line) and time-shuffled (dashed line) data. Note the significant decrease in PLV between 2-5 Hz in MS pups. (ii) Same as C. (i) for female control (n= 14) and MS (n=13) pups. D. (i) Average PLV spectra between activity in PL and BLA of control male (straight line) and female pups (dotted line) for original data and time-shuffled (dashed lines) data. (ii) Same as D. (i) for MS male and female pups. Note the significant decrease in PLV between 2-5 Hz in MS male versus female pups. In A. (iii), B. (iii), C. and D., the shaded areas correspond to S.E.M. * p<0.05.

In summary, this suggests that MS exposure specifically impairs the coupling synchrony between PL and BLA in delta/low theta range in males whereas synchrony in females remains unaffected.

## Discussion

It is well established that ELS results in long-lasting changes in the function of the cortico-limbic circuitry, underlying emotional and social behavior. Nevertheless, how exactly ELS affects the emerging projections from mPFC to BLA and how these early changes contribute to the enduring dysfunction of the circuitry remains poorly understood. In this study, we report in MS treated male rats, aberrant mPFC inputs to BLA GABAergic interneurons transiently shift the E/I balance towards enhanced inhibition, which raises the LTP induction threshold in cortico-amygdaloid synapses. The enhanced GABAergic activity after MS exposure in males associates with lower synchronization between mPFC and BLA already at P14-15. Impaired plasticity and synchronization during the sensitive period of circuit refinement may contribute to long-lasting functional changes in the cortico-amygdaloid circuitry that predispose to neuropsychological conditions later on in life.

### Sex-specific effects of ELS on mPFC-BLA anatomical connectivity

Consistent with several previous studies (reviewed by Murthy and Gould, 2018), MS resulted in an increase in anxiety-like behaviors in male rats but had no significant effects on females. Likewise, the observed effects of MS on mPFC-BLA connectivity were mostly male-specific. As an exception, inter-hemispheric innervation from the mPFC to the contralateral amygdala was significantly lower in female rats exposed to MS as compared to controls. In control rats, mPFC-BLA projections developed earlier in females as compared to males, with the ipsilateral innervation emerging earlier as compared to contralateral one. Therefore, our data supports the hypothesis that the timing of the stress exposure relative to the developmental stage is critical in defining the sex – specific outcome (van Bodegom et al., 2017).

Differences in cortico-amygdaloid connectivity have been previously shown in rodents exposed to ELS. Similar to our data, ELS is reported to increase or have no effect on structural connectivity between mPFC and BLA in males (Bolton et al, 2018; Manzano Nieves et al., 2020; White et al., 2020) and decrease functional connectivity (Yan et al., 2017; Guadagno et al., 2018; Honeycutt et al., 2020). Interestingly, in contrast to MS, the limited bedding and nesting (LBN) model of ELS (Walker et al., 2017) predisposes to adult stage anxiety-like behaviors also in females (Demaestri et al., 2020) and associates with a developmentally transient increase in PL-BLA anatomical connectivity (Manzano Nieves et al., 2020). Thus, developmental increase in mPFC-BLA structural connectivity in the context of ELS strongly correlates with emergence of anxiety-like behaviors later on in life in both sexes. However, the cell types involved in the hyper-connectivity have not been identified before. Here, we show for the first time that ELS increases mPFC projections particularly to GABAergic neurons in the BLA.

### ELS results in a developmentally transient increase in GABAergic activity in the amygdala

E/I balance within the mPFC-BLA pathway is shifted towards inhibition during adolescence in rodents (Arruda Carvalho et al., 2017), yet how this development is affected by ELS has not been studied previously. In order to understand the functional consequences of the aberrant mPFC innervation on BLA circuitry, we first recorded ongoing spontaneous synaptic activity from LA and BA principal neurons. MS had no effect on glutamatergic transmission but enhanced the frequency of GABAergic events in LA at the time (P14-15) when the transient increase in anatomical connectivity between mPFC and BLA was observed in males. Mechanistically, this effect can be fully explained by an ELS dependent increase in excitatory input from mPFC to BLA GABAergic neurons. First, the differences in GABAergic synaptic transmission between control and MS rats depended on intact glutamatergic transmission. Second, tracing data indicated that the relative proportion of mPFC inputs targeting GABAergic neurons in BLA was higher in MS exposed rats as compared to controls. Finally, the strength of feedforward inhibition (FFI) in response to selective optogenetic activation of mPFC inputs was enhanced after MS both in LA and BA. These data support that during development, the number of direct mPFC inputs to BLA interneurons is a critical stress-sensitive factor controlling the recruitment of GABAergic microcircuits and E/I balance in the BLA in response to mPFC activation.

Ionotropic glutamate receptor-mediated activity is required to regulate the subtype specific development of neuronal morphology in GABAergic interneurons (De Marco García et al., 2011). Therefore, changes in the glutamatergic connectivity could drive alterations in the morphological and neurochemical maturation of the interneurons during ELS and result in secondary changes that further influence E/I balance. Our RT-qPCR analysis did not reveal significant effects of MS on expression of molecules implicated in maturation of GABAergic neurons at P14-15, but the density of BLA PV+ neurons was lower after MS as compared to controls. However, previous work has reported significant changes in density, neurochemistry and morphology of interneurons after ELS, both in adult as well as in adolescent BLA (e.g. Gilabert-Juan et al., 2011; Castillo-Gómez et al., 2017; Giachino et al., 2007; Seidel et al., 2008; Santiago et al., 2018; Manzano Nieves et al., 2020). Receptors for stress-related hormones are expressed in various different types of neurons (Hartmann et al., 2017; Calakos et al., 2017; McKlveen et al., 2016). Experiments using mice with cell type-specific knockout of glucocorticoid receptors have found that forebrain glutamatergic, but not GABAergic, neurons drive the circuit changes mediating fear and anxiety behaviors after stress in adults (Hartmann et al. 2017). Similar studies in the context of ELS will help to reveal whether ELS affects maturation of interneurons directly or indirectly, via alterations on glutamatergic inputs.

### Enhanced GABAergic activity after ELS attenuates LTP induction in the cortico-amygdaloid pathway

In adult BLA, susceptibility of glutamatergic synapses to LTP is tightly controlled by GABAergic inhibition of principal neurons (Watanabe et al., 1995; Bissiere & Lüthi 2003; Shin et al., 2006; Bazelot et al., 2015). Furthermore, prior stress modulates amygdala LTP in adult (Vouimba et al., 2004; Rodríguez Manzanares et al., 2005; Suvrathan et al., 2013) and juvenile rats (Danielewicz and Hess, 2014; Guadagno et al., 2020). Our data expands these findings to show that the transiently increased GABAergic drive after ELS is sufficient to increase the threshold for LTP induction in the LA already at P18 – P21, coinciding with the rapid development and refinement of cortico-amygdaloid connectivity.

Interestingly, in the presence of GABA-A receptor antagonists, the level of synaptic potentiation in response to a pairing protocol was higher in MS males as compared to controls. One possibility to explain these data is that ELS generates functionally silent synapses, similar to that observed in the adult BLA after chronic stress and observational fear paradigms (Suvrathan et al., 2014; Ito et al., 2015). Silent synapses create a substrate permissive for plasticity, without affecting basal strength of synaptic transmission (Kerchner and Nicoll, 2008). Consistent with this idea, previous work has identified that BLA neurons in preweaning (P20) but not adult males that have experienced ELS show an increase in dendritic length and number of spines (Guadagno et al., 2018; but see Krugers et al., 2012), yet no associated changes in the mEPSCs, reflecting the density of functional inputs (Guadagno et al., 2018).

### Early effects of ELS on mPFC-BLA functional connectivity

Abnormal prefrontal-amygdala functional connectivity is one of the hallmarks of anxiety disorders in humans with a history of ELS (VanTieghem and Tottenham, 2018) and has also previously been observed in animal models of ELS using resting-state functional MRI (Yan et al., 2017; Guadagno et al., 2018; Honeycutt et al., 2020). Our data with simultaneous LFP recordings from mPFC and BLA demonstrates impaired synchronization of mPFC and BLA activities at the 2-5 Hz frequency range immediately after ELS exposure (P14-15) in males. Consistent with early dysfunction of the cortico-amygdaloid circuitry, aberrant social behavior in ELS exposed pups can be observed as early as P13 (Raineki et al., 2019) and deficits in fear recall at P21 (Manzano Nieves et al., 2020).

In the adult, mPFC-BLA theta range synchronization during fear and anxiety behavior is mainly driven by mPFC activity, entraining the firing of local BLA neurons to mPFC rhythms (Karalis et al., 2016, Likhtik et al., 2014). Similar to hippocampus, however, the local interneurons in the BLA likely have a critical role in generation of rhythmic activity by gating the firing of principal cells (Buzsaki, 2002). The mistargeting of the mPFC axons and subsequent shift in the BLA E/I balance could thus represent one mechanism behind the deficient mPFC –BLA functional coupling after ELS in young males.

During long-range synchrony, the excitability of BLA neurons is modulated by oscillations in other structures, which could play an important role in plasticity induction *in vivo* (Bocchio et al., 2017). Thus, stress-dependent early alterations in the excitability and synchronization of the cortico-amygdaloid circuitry could have enduring effects on cortico-limbic reactivity, via perturbing activity-dependent synaptic refinement.

## MATERIALS AND METHODS

### Animals

Experiments were performed using male and female Wistar rats. All experiments were done in accordance with the University of Helsinki Animal Welfare Guidelines and approved by the Animal Experiment Board in Finland.

### Maternal separation

The maternal separation protocol was done essentially as described (Englund et al., 2021). Briefly, pups were assigned to two experimental groups randomly on postnatal day (P)2. In the MS group, the pups and the dam were separated daily for 180 min period from P2 until P14 (MS group), while littermate controls remained with the dam in their home cage. Heat pads were used to maintain body temperature during the separation. On postnatal day 21, the litters were weaned and the animals were maintained under standard housing conditions (2–5 animals of the same sex/cage) until adulthood. The animals were sacrificed for experiments between 9 a.m. and 3 p.m., during the light period (lights on 6 a.m., off 6 p.m.). The stage of the estrous cycle was not controlled. Behavioral testing was done at P60 – 80 as described previously (Englund et al., 2021).

### Viral injections

The following AAV viral vectors were used: anterograde tracer pAAV-hSyn-EGFP (Addgene 50465 –AAV1) and channelrhodopsin pAAV-CaMKIIa-hChR2(H134R)-EYFP (Addgene 26969-AAV1). The viral particles were injected to the mPFC of neonatal (P4-7) or adolescent (P30) rat mPFC under isoflurane anesthesia essentially as described (Ryazantseva et al., 2020). Briefly, small burr holes were drilled above the region of interest, after which 100-150 nl of viral suspension was injected at two depths, targeting the prelimbic and infralimbic areas of the prefrontal cortex. After each injection, the pipette was held in place for 5 min before moving on. For optogenetics, viral particles were injected bilaterally at P4-7, using the following coordinates: 2.2 (AP), ±0.25 (ML), −1/-2 (DV) and 2.8 (AP), ±0.25 (ML), −1/-2 (DV). For tracing, injections were made to one hemisphere only (coordinates P4-6: 2.2 (AP), 0.25 (ML), −1/-2 (DV) and 2.8 (AP), 0.25 (ML), −1/-2 (DV); P30: 2(AP), 0.5(ML), −1.5/-2.5 (DV) and 2.5 (AP), 0.5(ML), −1.5/-2.5 (DV)). Once the injections were completed, animal was allowed to wake up and recover on the heat pad for 10 min, prior to placing them back into their home cage. Animal welfare was observed daily after the procedure.

### Visualization of the viral tracers

The rats were perfused transcardially with PBS and 4% PFA under deep isoflurane anesthesia. The brains were removed and postfixed in 4% PFA overnight at 4 °C, after which the brains were immersed in 30% sucrose/PBS solution for several days before transferring to cryoprotectant (Tissue–Tek, Sakura). Coronal cryosections (20 μm) were cut using Leica CM3050 cryotome and stored at −20 °C. Immunohistochemical stainings were performed in some of the sections. After washing with PBS-T (PBS + 0.2% Triton X-100), the cryosections were briefly treated with 0.5% SDS (5 min), washed again and incubated 30min at RT in blocking solution (1% BSA + 4% NGS in PBS). The primary antibody (Glutamic acid decarboxylase isoform 67, mouse; DMEM Millipore MAB5406; 1:500) was added to PBS-T and incubated 48 h at 4 °C. After washing with PBS, the sections were placed in the solution containing the secondary antibody (anti-mouse Alexa Fluor 588, Life Technologies; 1:200 in PBS-T for 2h RT). All sections were rinsed with PBS before mounting them in Vectashield with DAPI (Vector Laboratories). Images were taken with a Zeiss^®^ AX10 microscope and further analyzed with ImageJ® and Zen2.

### RT-qPCR and Immunohistochemistry of GABAergic markers

BLA was dissected from 400 μm thick vibratome sections cut from the brain of control and MS Wistar rats at postnatal day (P)14 or P50. Purification of total RNA, cDNA synthesis and the real-time quantitative PCR was carried out essentially as described (Ryazantseva et al., 2020), using the following probes:

*Pvalb* F: 5′-TTCTGGACAAAGACAAAAGTGG-3′ R: 5′-CTGAGGAGAAGCCCTTCAGA-3′; *Slc12a2* F: 5′-TCCTCAGTCAGCCATACCCAAA-3′ R: 5′-TCCCGGACAACACAAGAACCT-3′;
*Slc12a5* F: 5′-TCCTCAAACAGATGCACCTCACCA-3′ R: 5′-ACGCTGTCTCTTCGGGAACATTGA-3′
*SST F:* 5′-GCCCAACCAGACAGAGAACGATGC-3′ R: 5′-GCTGGGTTCGAGTTGGCAGACCTC-3′.

For immunohistochemical stainings, mice were intracardially perfused with 4% PFA, the brains were removed and postfixed in 4% PFA. All samples were dehydrated and embedded in paraffin using automated Leica tissue processor, and sectioned at 5 μm. After deparaffination, the antigen retrieval was done by heating the sections in 10 mM sodium-citrate buffer, pH 6.0, in a microwave oven (10 min). The sections were washed with TBS and blocked for 30–60 min solution containing TBS + 10% NGS +3%BSA+ 0.25 % Triton X-100. The primary antibodies (Anti-Parvalbumin (195 004, Synaptic Systems) 1:2000, Anti Anti-Somatostatin (SAB4502861, Sigma-Aldrich) 1:100) were added to the blocking solution and incubated overnight at RT. After rinsing with TBS, the sections were placed in the secondary antibody (Alexa Fluor 488/594-conjugated goat anti-guinea pig /rabbit secondary antibodies, Invitrogen) in TBS + 0.1% Triton X-100 for 3 h at RT. Sections were rinsed with TBS before mounting them in Vectashield with DAPI (Vector). Immunostaining was visualized with an Olympus BX61 microscope and counting of cell density was done using ImageJ software.

### In vitro Electrophysiology

#### Preparation of acute slices

P14-21 rat pups were anesthetized with isoflurane and quickly decapitated. The brain was extracted and immediately placed in carbogenated (95% O_2_/5%CO_2_) ice-cold sucrose-based dissection solution containing (in mM): 87 NaCl, 25 NaHCO3, 2.5 KCl, 1.25 NaH2PO4, 7 MgSO4, 0.5 CaCl2, 25 D-glucose, 50 sucrose. The cerebellum and a small part of the prefrontal cortex were trimmed off and the remaining block was glued to a stage and transferred to a vibratome (Leica VT 1200S) to obtain 350 μm thick brain slices. Slices containing the amygdala were transferred into a slice holder in a heat bath (37 C) containing standard ACSF with added 1 mM MgSO4. After 30 minutes in the heat bath, the slices were stored in room temperature. Slices from adult (P50) rats were prepared using NMDG-based solution as described (Englund et al., 2021).

#### Electrophysiological recordings

After 1-5h of recovery, the slices were placed in a submerged heated (32-34°C) recording chamber and perfused with standard ACSF containing (in mM): 124 NaCl, 3 KCl, 1.25 NaH_2_PO_4_, 26 NaHCO_3_, 15 glucose 1 MgSO_4_ · 7H_2_O, 2 CaCl_2_ at the speed of 1-2 ml/minute. Whole-cell patch clamp recordings were done from LA and BA neurons under visual guidance using patch electrodes with resistance of 3–7 MΩ. Cells with capacitance > 50 pF and spike frequency adaptation to membrane depolarization were considered principal neurons. Multiclamp 700B amplifier (Molecular Devices), Digidata 1322 (Molecular Devices) or NI USB-6341 A/D board (National Instruments) and WinLTP version 2.20 (Anderson and Collingridge, 2007) or pClamp 11.0 software were used for data collection, with low pass filter (10 kHz) and sampling rate of 10-20 kHz. In all voltage clamp recordings, uncompensated series resistance (Rs) was monitored by measuring the peak amplitude of the fast whole-cell capacitance current in response to a 5 mV step. Only experiments where Rs < 30 MΩ, and with < 20 % change in Rs during the experiment, were included in analysis. The drugs were purchased from Hello Bio (Picrotoxin, D-(-)-2-amino-5-phosphonopentanoic acid (D-AP5), and CNQX).

#### Spontaneous glutamatergic and GABAergic synaptic responses (sEPSCs and sIPSCs)

were recorded from LA neurons at a holding potential of −30 mV using electrodes filled with a potassium based low chloride filling solution containing (in mM): 135 K-gluconate, 10 HEPES, 2 KCl, 2 CaOH2, 2 EGTA, 4 Mg-ATP, 0.5 Na-GTP. The membrane potential was first clamped at −70 mV, and the cell was allowed to equilibrate with the filling solution for 5 min, after which the membrane potential was slowly raised to −30 mV in order to record both sEPSC and sIPSC simultaneously.

#### Spontaneous pharmacologically isolated sIPSCs

were recorded using a cesium-based filling solution (pH 7.2, 280 mOsm) containing (in mM): 145 mM CsCl, 10 mM HEPES, 5 mM EGTA, 2 mM MgCl2, 2 mM CaCl2, 4 mM ATP, 0.33 mM GTP. CNQX (10 μM) and D-AP5 (50 μM) were included in the perfusion solution to antagonize AMPAR and NMDAR-mediated components of synaptic transmission, respectively. The membrane potential was clamped at −70 mV, and the cell was allowed to equilibrate with the filling solution for 5–10 min prior to recording sIPSCs.

#### The strength of feedforward inhibition

was recorded using a cesium-based filling solution (pH 7.2, 280 mOsm) containing (in mM): 125.5 Cs-methanesulfonate, 10 HEPES, 5 EGTA, 8 NaCl, 5 QX314, 4 Mg-ATP, 0.3 Na-GTP.

The membrane potential was first clamped at −70 mV, and the cell was allowed to equilibrate with the filling solution for 5-10 minutes. EPSCs were evoked by light stimulation (pulse width: 1 ms, intensity: 4 - 30 mW), after which the membrane potential was slowly raised to 0 mV, the reversal potential of glutamatergic currents, in order to isolate fast GABAA-R mediated currents. In these experiments, the Vm values were corrected for the calculated LJP. The cell was allowed to stabilize at 0 mV for 5-10 minutes prior to baseline measurement. 10 μM CNQX was applied at the end of the experiments to ensure that the recorded IPSC was disynaptic (> 80% block of the IPSC amplitude).

#### Field LTP recordings

In order to obtain stable recordings, slices were allowed to sit in the recording chamber for 15 – 30 minutes before the experiment was started. fEPSPs were evoked with a bipolar stimulation electrodes (nickel-chromium wire) placed in the external capsule to mimic activation of cortical inputs, while the recording electrode (resistance 1-3 MΩ, filled with standard ACSF) was placed in the lateral amygdala (LA). After a stable baseline, LTP was induced using either high frequency stimulation (100Hz/s) or thetaburst stimulation (6 bursts of 8/100Hz at 5Hz), repeated 3 times with 5 min intervals.

#### Pairing-induced LTP

were recorded using a potassium-based filling solution (pH 7.2, 278 mOsm) containing (in mM): 135 K-gluconate, 10 HEPES, 0.2 EGTA, 8 KCl, 4 Mg-ATP, 0.3 Na-GTP. Picrotoxin (100 μM) was included in the perfusion solution to antagonize fast GABAA mediated components of synaptic transmission. The membrane potential was clamped at −70 mV to obtain a stable EPSC baseline. LTP was induced by a pairing protocol by briefly depolarizing the cell to +10 mV and applying a single train of theta burst stimulation (6 x 8/100Hz at 5Hz), after which the cell was clamped back down to −70 mV.

#### Data analysis

WinLTP program was used to calculate the peak amplitude of the evoked synaptic responses. For EPSC/dIPSC ratios, 7-10 responses were averaged in each experimental condition. EPSC/dIPSC ratio was calculated as the amplitude ratio of response at −70 mV/response at 0 mV. The frequency and amplitude of spontaneous synaptic events was analyzed using Minianalysis program 6.0.3. sIPSCs and sEPSCs were identified in the analysis as outward and inward currents, respectively, with typical kinetics, that were at least 3 times the amplitude of the baseline level of noise. Averages for baseline, drug application and washout were calculated over a 10 minute period. n numbers for electrophysiological data are given as the number of cells /slices, followed by number of animals in parenthesis.

### In vivo electrophysiology

#### Surgical preparation

Extracellular recordings were performed in medial prefrontal cortex (mPFC) and basolateral amygdala (BLA) of postnatal day (P)14-15 male and female rats.

Anaesthesia was induced with isoflurane (4%) followed by i.p. administration of urethane (1g/kg; Sigma-Aldrich). Isoflurane administration (1-2%) continued during surgery to ensure deep anaesthesia. Small burr holes were drilled above the regions of interest (PFC: 1.2-1.5 mm anterior to bregma and 0.3 mm from the midline, BLA: base of the rhinal fissure) and the bone was carefully removed. The head of the pup was fixed into the stereotaxic apparatus (Stoelting, Wood Dale, IL) using two metal bars fixed with dental cement on the nasal and occipital bones, respectively. The body temperature was maintained at 37 °C with a thermoregulated heating blanket (Supertech Instruments, UK). A urethane top-up of 1/3 of the original dose was administered if needed. Multielectrode arrays (Silicon Michigan probes, NeuroNexus Technologies) were inserted perpendicularly to the skull surface into PFC until a depth of 3.5-4 mm and at 40° from the vertical plane into BLA at a depth of 4.5 mm. The electrodes were labeled with DiI (1,1’-dioctadecyl-3,3,3’,3’-tetramethyl indocarbocyanine, Invitrogen) to enable reconstruction of the electrode tracks post mortem (Figure 6A (i), B (i)). Two silver wires were inserted into the cerebellum and served as ground and reference electrodes.

#### Recording protocol

After 15-20 min recovery period, simultaneous recordings of local field potential (LFP) and multi-unit activity (MUA) were performed from the prelimbic subdivision (PL) of mPFC and basolateral amygdala (BLA) using 32-channel four shank (4×8) Michigan electrodes (0.5-3 MΩ). The distance between shanks was 200 μm and the recording sites were separated by 100 or 200 μm. The position of recording sites over the PL and BLA area was confirmed by post-mortem histological evaluation. Both LFP and MUA were recorded for at least 45 min at a sampling rate of 32 or 20 kHz using a multi-channel extracellular amplifier (Digital Lynx 4SX, Neuralynx or Smartbox, Neuronexus) and the corresponding acquisition software. During recording the signal was band-pass filtered between 0.1 Hz and 8 kHz (Neuralynx) or 0.001 Hz and 5 kHz (Neuronexus).

#### Data analysis

Channels for analysis were selected on the basis of post-mortem histological investigation, i.e. which recording sites of fluorescently-marked electrodes were confined to PL and BLA (n= 55 animals in total). Here, recordings from individual channels with best signal-to-noise ratio over superficial layers II/II of PL and medial aspects of predominantly anterior basal amygdala were analysed according to the anatomical development of this projection (Arruda-Carvalho et al. 2017). In total four animals were excluded due to visible heartbeat artifacts in the recordings. Data were imported and analyzed off-line using custom-written tools in Matlab software version 2020b (Mathworks, Natick, MA). The signals were low-pass filtered (<1500 Hz) using a third-order butterworth filter before downsampling to 2000 Hz. An IIR notch filter was applied at 50 Hz (q-factor = 35) using the Matlab function *filtfilt* to avoid phase distortions. All data analysis was performed blind without knowing the group belonging of the pup.

For spectral analysis, a continuous wavelet transform was performed on the signals. The LFP signal was convolved with a series of complex Morlet wavelets with the length of 7 cycles and 50 center frequencies logarithmically distributed in the band 1-100 Hz. For visualization of the trace spectrogram (see wavelet in

Fig. X A (ii), B(ii)), 100 center frequencies were linearly distributed in the same band. Spectral power was defined as the square of the absolute value of wavelet transformed LFP converted to decibel.

Phase relationships between amygdala and mPFC signals were analyzed by phase locking value (PLV), which is defined as 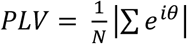, where θ is the difference between the instantaneous phase of the signals. For wavelet transformed LFPs X and Y, PLV was calculated as 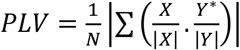,, where N is the length of the signal, Y* - complex conjugate of Y.

Confidence limits for PLV were determined by Monte Carlo simulation. For this, 3 s-long LFP segments from one region were shuffled and PLV calculated between the shuffled LFP from one region and the original LFP from the other region. After 100 iterations, the 95^th^ percentile of the resulting distribution was used as a significance threshold.

### Statistical analysis

All statistical analyses were done on raw (not normalized) data using Sigma Plot or Prism GraphPad software. The sample size was based on previous experience on similar experiments. All data were first assessed for normality and homogeneity of variance and the statistical test was chosen accordingly. Differences between two groups were analyzed using two-tailed t-test or Mann–Whitney rank-sum test. Two-way ANOVA with Holm–Sidak post-hoc comparison was used to compare effects of sex and stress. For data that were not normally distributed, Mann–Whitney test was used for post-hoc pairwise comparison. The results were considered significant when *p* < 0.05. All the pooled data are given as mean ± SEM.

## Author Contributions

J.H., J.E., M.S.C. and T.A. conducted the maternal separation protocol and T.A. did the behavioral testing. J.H. and J.E. carried out the in vitro electrophysiological experiments and data analysis, J.H. performed all the experiments with optogenetics. Z.K. and H.H. did in vivo electrophysiological recordings and data analysis. S.P.A., K.K. and M.S.C. conducted the experiments with viral tracing and image analysis. A.S. did all the qPCR and immunohistochemical experiments with GABAergic markers. S.E.L., H.H. and T.T. provided resources for the experimental work and supervised the project. S.E.L conceptualized and coordinated the project. The manuscript was written by S.E.L. and J.H. with significant contributions from all authors.

## Acknowledgements

We thank Kirsi Kolehmainen and the personnel in the Finnish Centre for Laboratory Animal Pathology (FCLAP) for expert technical help. This study was financially supported by the Academy of Finland, Sigrid Juselius Foundation, Finnish Cultural Foundation and EduFI.

---

**Suppl Figure 1.**
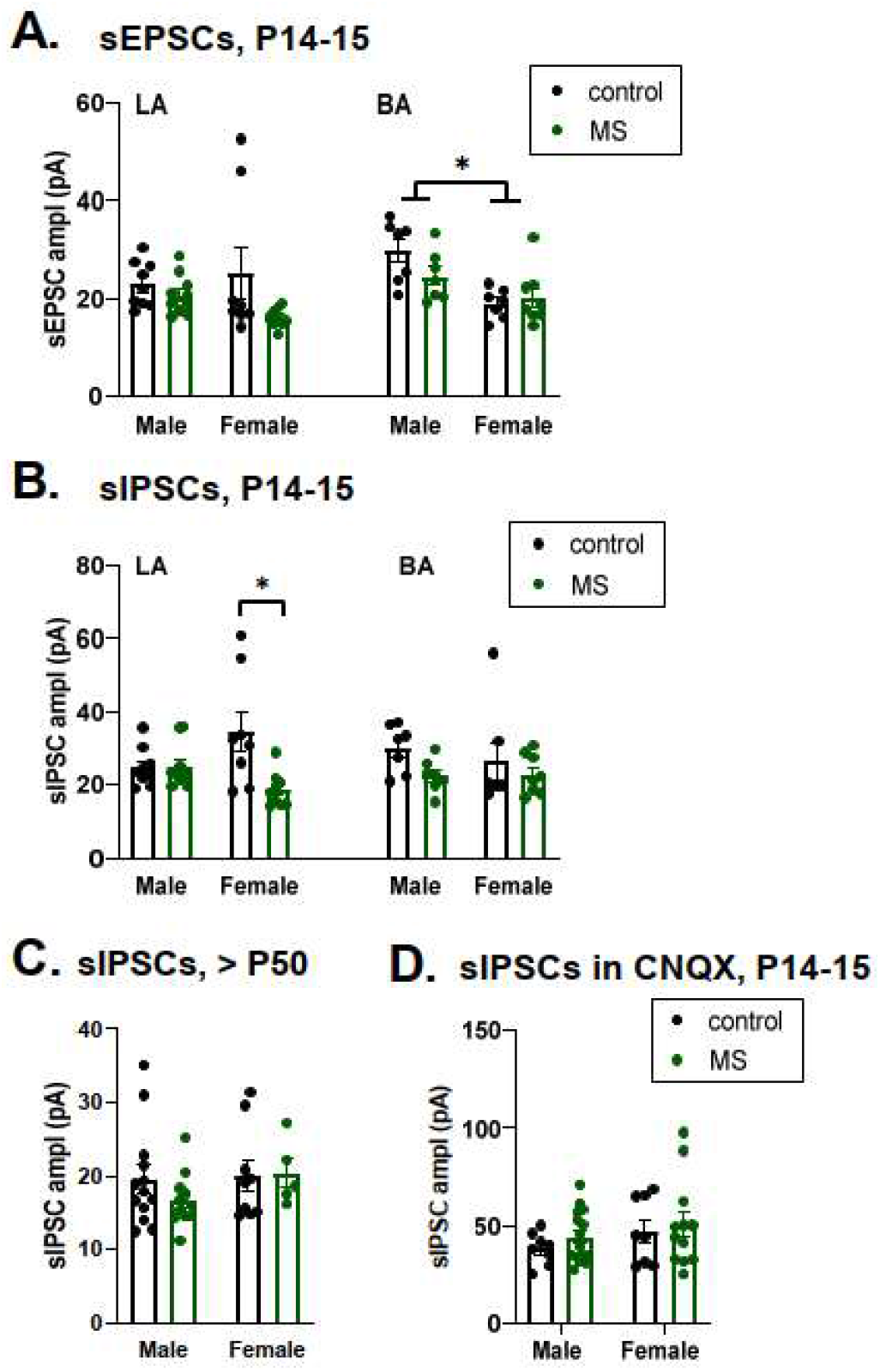
MS is associated with developmentally restricted and sex specific changes in GABAergic activity in the amygdala. Data on the amplitude of the spontaneous events, for the same recordings that are shown in the figure 3. A. Pooled data on the amplitude of sEPSCs in recordings from LA and BA neurons in control (black symbols) and MS (green symbols) juvenile (P14-15) rats [control male n= 9(4 animals), female n=8(3), MS male n=10 (4), female n=9(3)]. In BA, sEPSC amplitude was significantly affected by sex (F_(1, 25)_ = 15.24, p=0.0006). B. Pooled data on the amplitude of sIPSCs, from the same cells as in A. sIPSC amplitude in LA was significantly lower in MS females compared to controls (* p<0.05; one –way ANOVA on ranks). C. The amplitude of sIPSC in LA neurons from adult control and MS treated rats [control male n=12(10), female n=9(7), MS male n=10 (8), MS female n=5(3)]. No significant differences between groups were detected. D. Pooled data on the amplitude of sIPSCs in LA neurons from juvenile (P14-15) control and MS rats, recorded in the presence of CNQX to block glutamatergic activity. These recordings were done using high chloride concentration in the pipette filling solution, which improves detection of small GABAergic events that are now observed as inward currents. Under these conditions, no significant differences in sIPSCs were detected between the groups [control male, n=9(3), female n=8(3); MS male, n=16 (4), female n=13(4)].

**Suppl Figure 2.**
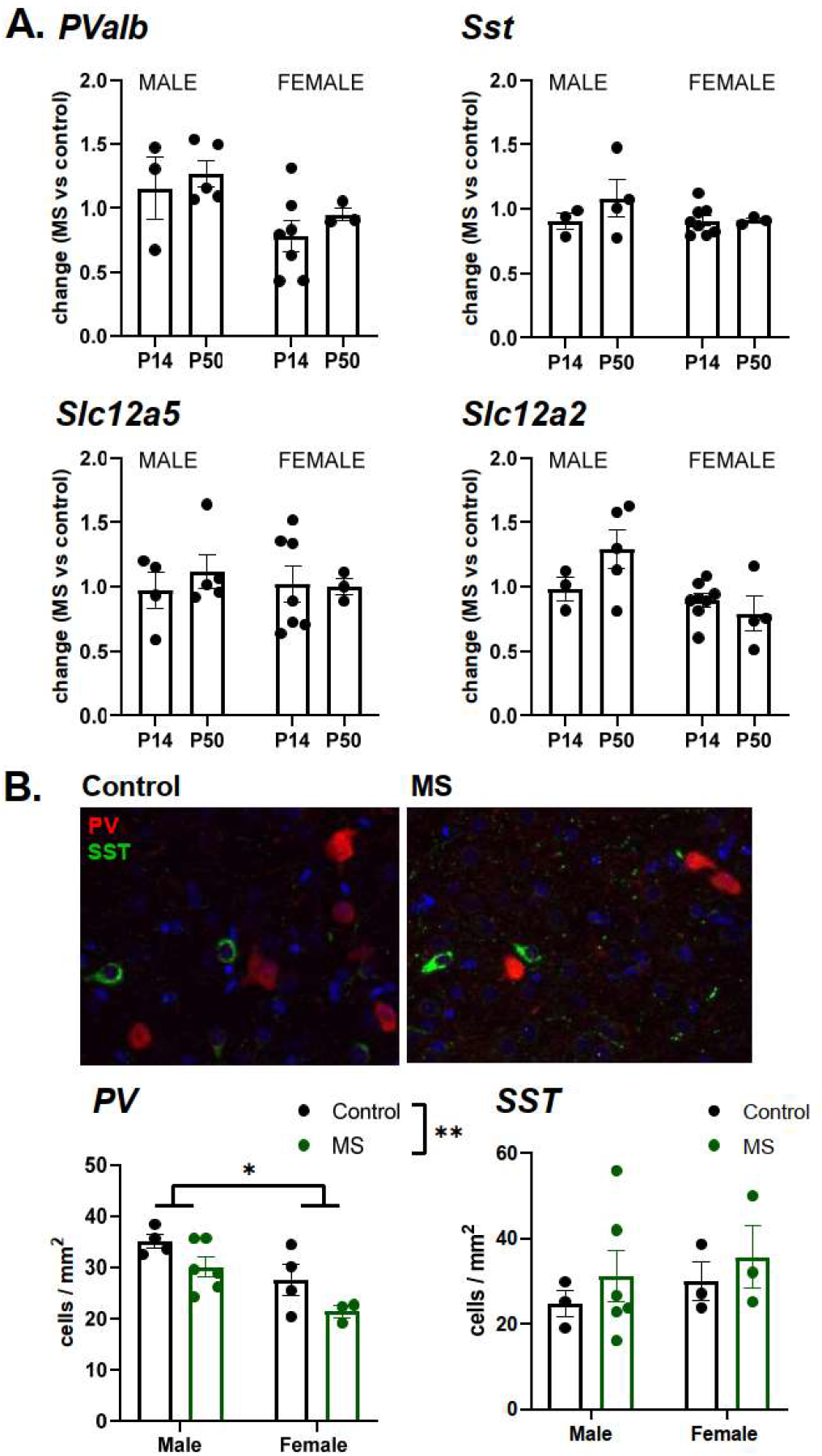
Effect of MS on expression of GABAergic markers in the BLA. A. RT-qPCR analysis of *PValb, Sst, Slc12a5 and Slc12a2* mRNA expression levels in the basolateral amygdala (BLA) in MS and control rats. The pooled data shows the level of gene expression in MS animals, normalized to controls. n=3-8 rats /group (P14 Male n=3-4, Female n=7-8; P50 Male n=4-5, Female n=3-4). The MS treatment had no significant effect on expression of any of the mRNAs studied. B. Example images of immunohistological staining for parvalbumin (PV, red) and somatostatin (SST, green) in basolateral amygdala (P14), in male control and MS rats. Pooled data shows the average density of PV+ and SST+ neurons in male and female control and MS rats. PV cell density was significantly lower in females as compared to males (F_(1,13)_ = 6.573, p=0.02) and lower in MS group as compared to controls (F_(1,13)_ = 13.40, p=0.03). No significant differences were detected in the density of SST+ neurons. n=3-6 rats / group (control male n=3-4, female n=3-4; MS male n=6, female n=3).

**Suppl. Figure 3.**
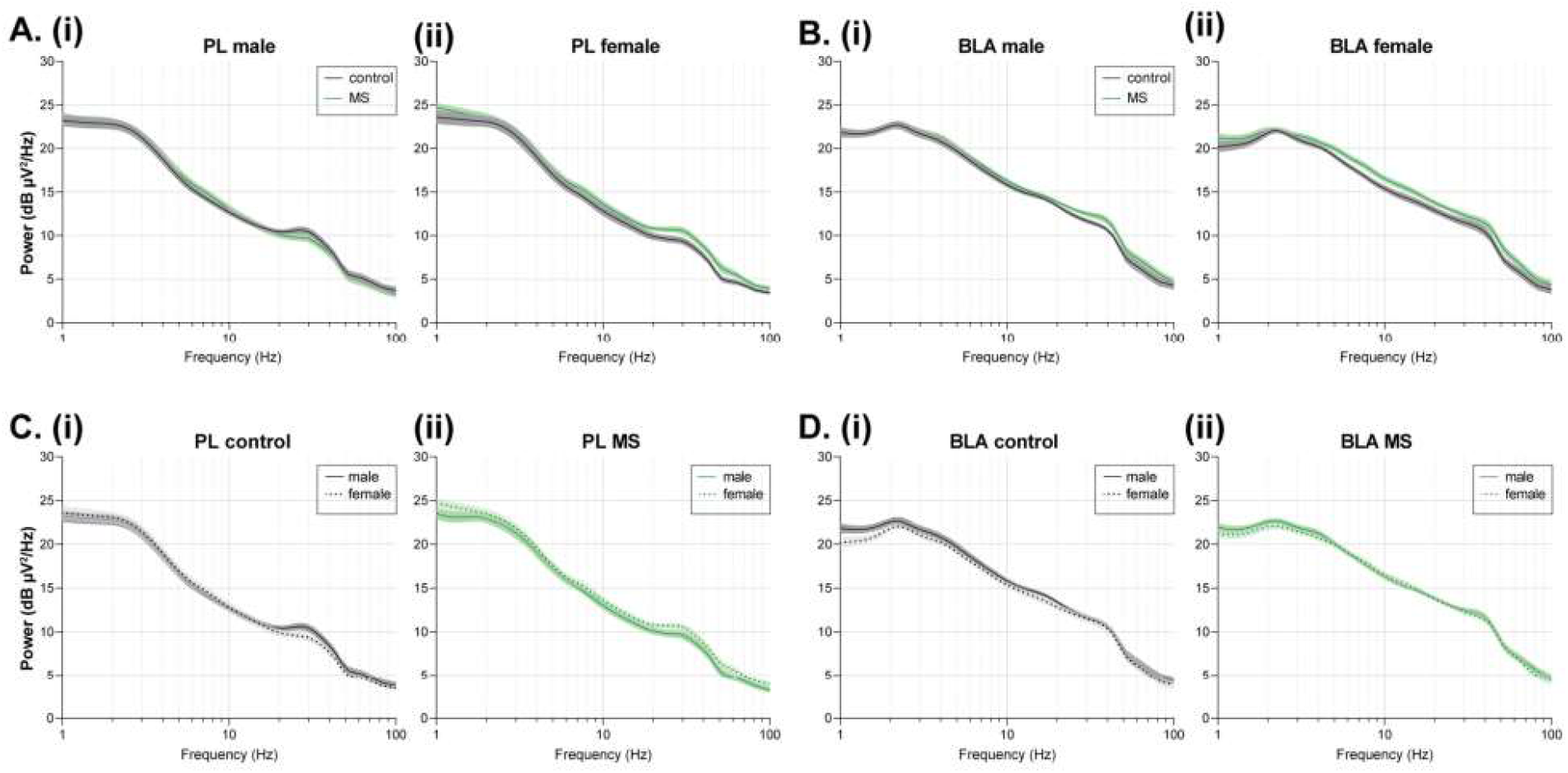
MS does not affect oscillatory power neither in PL nor in BLA. A. (i) Average power spectra of oscillatory activity in PL for male control (n=11, black) and MS (n= 13, green) rat pups. (ii) Same as A. (i) for female control (n=14) and MS (n=13) pups. B. (i) Average power spectra of oscillatory activity in BLA for male control (n=11, black) and MS (n=13, green) rat pups. (ii) Same as B. (i) for female control (n=14) and MS (n=13) pups. C. (i) Average power spectra of oscillatory activity in PL for control male (straight line) and female (dotted line) rat pups. (ii) Same as C. (i) for MS male and female pups. D. (i) Average power spectra of oscillatory activity in BLA for control male (straight line) and female (dotted line) rat pups. (ii) Same as D. (i) for MS male and female pups.

